# Marmosets adaptively accumulate dynamic sequential evidence to make decisions

**DOI:** 10.64898/2026.08.31.747938

**Authors:** Xianlin Xia, Li Bo, Ning-long Xu

## Abstract

Marmosets are a promising non-human primate model for studying the neural-circuit mechanisms of higher cognition. However, many aspects of their cognitive capacities remain to be characterized. Here, we developed a complex decision-making task in marmosets in which they accumulate dynamic sequential auditory evidence to make decisions. Behavioral analyses showed that the accumulation strategy dominated marmoset decision-making, and early evidence was weighted more heavily. Notably, late-evidence weights were relatively higher in difficult than in simple trials, an effect we term evidence-strength-dependent temporal weighting (ETW). After excluding other candidate behavioral mechanisms, we found that adaptive modulation of cognitive effort (e.g., attentional engagement with sensory evidence) across evidence strengths may be a key mechanism underlying this effect. A modified drift diffusion model (DDM), with diffusion noise parameterized separately for each evidence strength, reproduced the ETW effect. In the best-fitting model, diffusion noise negatively correlated with evidence strength, suggesting that cognitive effort may be upregulated as perceived task difficulty increases. Together, our work provides an important behavioral and computational foundation for unraveling neural-circuit mechanisms underlying complex decision-making in primates.

## Introduction

Common marmosets (*Callithrix jacchus*), as New World monkeys, are a promising non-human primate model in cognitive neuroscience(Magrou et al., 2024; Mitchell and Leopold, 2015). Compared with rhesus monkeys, their small body size and smooth cortex facilitate the application of neural circuit analysis tools such as two-photon imaging and optogenetics(Jendritza et al., 2023; Kondo et al., 2018). Additionally, their high reproductive rate enables rapid colony expansion, lowering experimental costs(Okano, 2021; Okano et al., 2012). In recent years, progress has been made in developing freely-moving or head-restrained behavioral paradigms in marmosets(Cloherty et al., 2020; Ebina et al., 2018; Nakamura et al., 2018; Sedaghat-Nejad et al., 2019). However, there remains a lack of complex cognitive behavioral paradigms for head-restrained marmosets because of the training difficulty of this species(Okano, 2021), which has limited their use in cognitive neuroscience over the past decades and many aspects of their cognitive capacities remain uncharacterized.

A hallmark of higher cognition is the capability of evaluating and accumulating multiple pieces of discrete evidence for decision-making. To study the behavioral strategies and the underlying neural mechanisms of this process, stimuli consisting of discrete sensory evidence are widely used in decision-making paradigms(Brunton et al., 2013; De Lange et al., 2010; Keung et al., 2019; Morcos and Harvey, 2016; Raposo et al., 2014; Tsunada et al., 2016; Waskom and Kiani, 2018; Wyart et al., 2012; Yates et al., 2017). For discrete evidence accumulation, a key question is how evidence is weighted over time. Studies across different species and behavioral paradigms show that the brain exhibits diverse temporal weighting patterns, including the primacy effect(Morcos and Harvey, 2016; Nienborg and Cumming, 2009; Yates et al., 2017), recency effect(Cheadle et al., 2014; Wyart et al., 2015), and uniform weighting (Brunton et al., 2013; Raposo et al., 2014; Wyart et al., 2012). The primacy effect is a dominant temporal weighting pattern in many non-human primate studies(Levi et al., 2023; Nienborg and Cumming, 2009; Yates et al., 2017), and the bounded accumulation mechanism is commonly used to account for this effect(Kiani et al., 2008; Ratcliff and McKoon, 2008). In this mechanism, a decision variable (DV) evolves between two decision bounds, driven by the input evidence and perturbed by random noise. Once the DV reaches either bound, the decision is terminated, and later evidence is ignored. A natural prediction of this mechanism is the ETW effect: when evidence strength is low, the DV takes longer to reach a bound, which allows later evidence to be incorporated more frequently and thus increases its contribution to the final decision. However, this prediction has not yet been directly tested, and it remains to be systematically evaluated whether the bounded accumulation mechanism alone is sufficient to account for the ETW effect.

Here, we developed a reliable decision-making task training framework for head-restrained marmosets, with which we established a novel sequential tone accumulation task in marmosets. Psychophysical weight analysis revealed the ETW effect in evidence accumulation. Computational modeling and behavioral analysis showed that neither the bounded accumulation mechanism alone nor post-stimulus memory sampling was sufficient to account for this effect. After excluding other candidate mechanisms, we found that adaptive modulation of cognitive effort across different evidence strengths may be a key mechanism underlying the ETW effect. Within the bounded accumulation model framework, cognitive effort can be captured by diffusion noise. A modified DDM in which diffusion noise was parameterized separately for each evidence strength successfully accounted for the ETW effect. The best-fitting model exhibited lower diffusion noise at weaker evidence strengths, suggesting that marmosets may adaptively upregulate internal cognitive effort as perceived task difficulty increases. Together, our work provides an important behavioral and computational foundation for investigating the neural-circuit mechanisms underlying complex decision-making in primates.

## Results

### Marmoset behavior training under head restraint

During the task training, a marmoset was held in a customized body tube and its head was restrained. The entire restraint device was positioned inside a custom-built soundproof chamber. We used two methods for head restriction. One was tight restraint (head fixation), in which a customized head plate was surgically implanted on the skull and the head plate was fixed by the holders (Figure 1A-D). The other was loose restraint, in which the animal’s neck was locked, and the head was restrained by a customized helmet (Figure 1E-H). Three spouts and a white LED were positioned in front of the marmoset. At the beginning of each trial, the white LED was turned on to guide the marmoset to lick the central spout for task initiation and was turned off following a successful initiation. The stimulus was then delivered after a short random delay (0.5-1 s). The marmoset was required to lick the left or right spout to report its decision and to obtain water reward.

**Figure 1.**
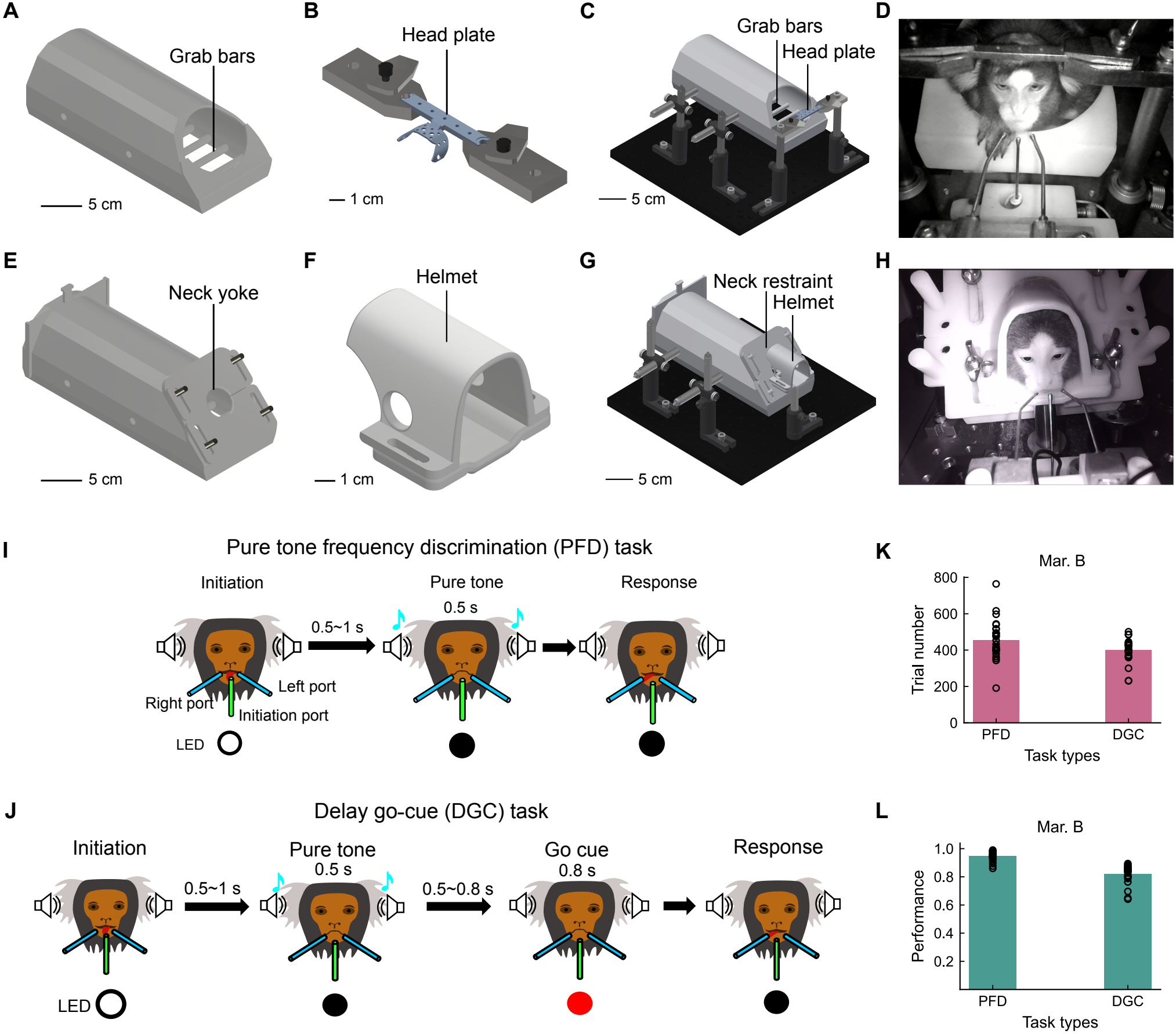
Head-restrained devices, behavioral paradigms and performance. (**A**) The head plate for head fixation. (**B**) The body restraint tube for head fixation. (**C**) The full view of head-fixation device. (**D**) A marmoset performs task under head fixation. (**E**) The body-restraint tube for loose head restraint. (**F**) The helmet for loose head restraint. (**G**) The full view of device for loose head restraint. (**H**) A marmoset performs task under loose head restraint. (**I**) Schematic of the pure tone frequency discrimination (PFD) task. Marmosets initiate trials by licking the central spout, guided by a front white LED. Successful initiation results in the white LED turning off. After a 0.5–1 s random delay, a pure tone is delivered. Marmosets lick the left or right spout to respond. (**J**) Schematic of the delayed go-cue (DGC) task. The task structure is similar to the PFD task. The key difference is that the marmosets cannot respond immediately after stimulus onset, but have to wait for a random delay (0.5–0.8 s). A red LED then signals marmosets to respond. (**K**) Mean number of trials completed by marmoset B in the PFD and DGC tasks; black circles indicate trial counts from individual sessions. (**L**) Mean performance of marmoset B in the PFD and DGC tasks; black circles indicate performance from individual sessions.

We designed two tasks to test the effectiveness of our training framework. Marmoset B sequentially learned the pure tone frequency discrimination (PFD) task (Figure 1I). and the delayed go-cue (DGC) task under head fixation (Figure 1J). For the first task, the marmoset was required to discriminate between the low-(4.5 kHz) and high-frequency (12.7 kHz) tones. To receive a reward, it needed to lick the left spout following the lower-frequency tone, and the right spout following the higher-frequency tone. The second task was an extension of the first, in which after stimulus onset, the marmoset was required to wait for a random delay period (0.5-0.8 s) before making a lick response. A red LED signaled the end of the delay period. In both tasks, Marmoset B completed 300 to 600 trials (average trial number was 454±21 for the PFD task and 400±12 for the DGC task, Figure 1K) in a single session and maintained performance above 80% across multiple training sessions (average accuracy for the PFD task was 94.61±0.62% and 82.09±1.3% for the DGC task, Figure 1L).

### DSTA task and behavioral performance

We further developed a sequential tone accumulation task in marmosets. In this task, the marmosets learned to discriminate the stimuli and report their decision after a go cue (Figure 2A). The stimulus consisted of a sequence of five discrete, equal-intensity pure tones, randomly alternating between high (11.8 kHz) and low (5.4 kHz) frequencies. Each tone lasted 0.125 s with an inter-tone interval of 0.125 s, yielding a total stimulus duration of 1.25 s. The marmosets were required to report which tone frequency was the majority in the tone sequence (Figure 2A).

**Figure 2.**
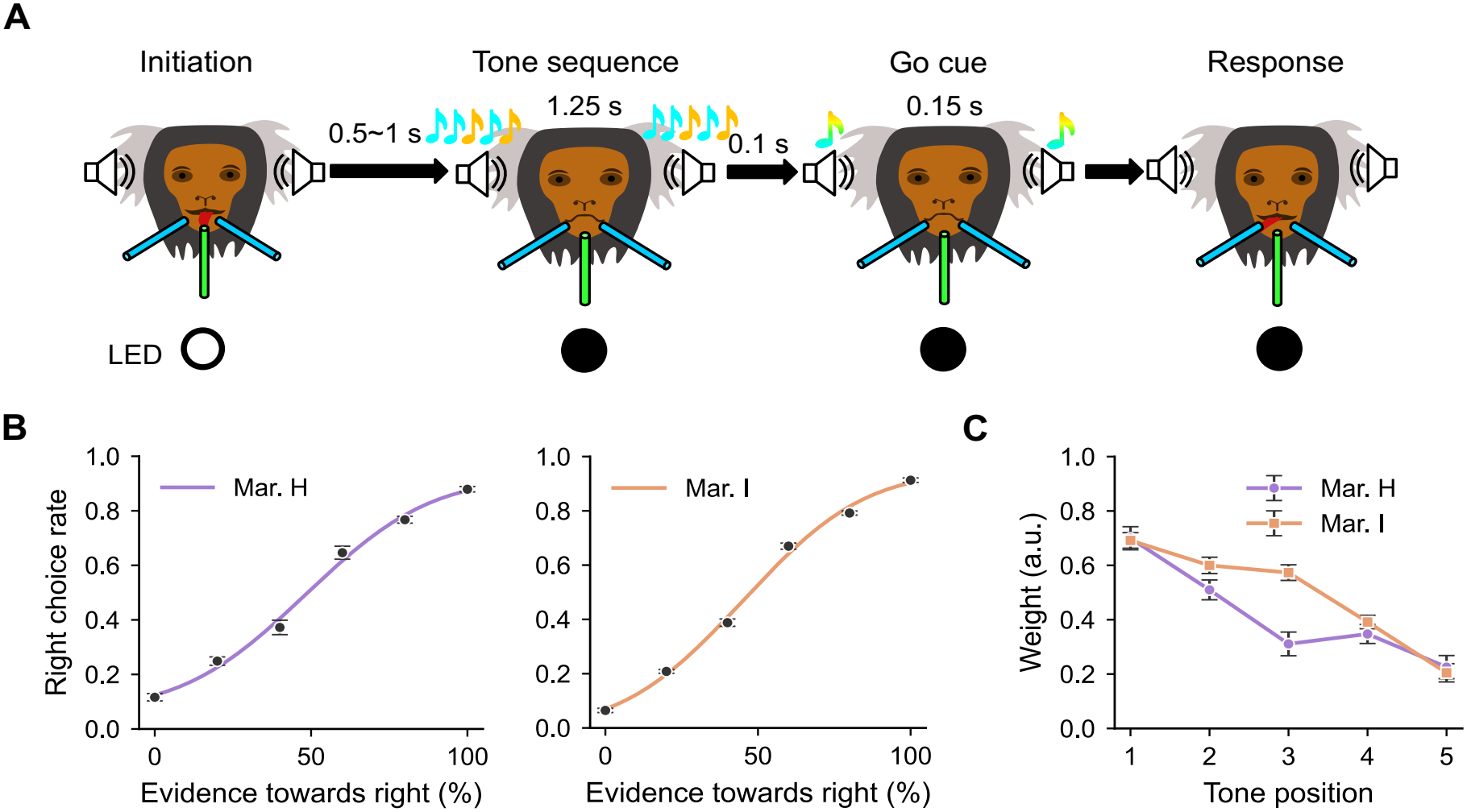
DSTA task design and behavioral performance. (**A**) Schematic of the dynamic sequential tone accumulation (DSTA) task. Marmosets initiate trials by licking the central spout, guided by a white LED. After a 0.5–1 s random delay, a 1.25 s tone sequence is delivered, followed by a short delay (0.1 s) and an auditory go cue (a 0.15 s sweep tone) signaling the response period. Marmosets are required to evaluate which frequency type predominates in the tone sequence and lick the left (more low-frequency tones) or right (more high-frequency tones) spout to report their decisions. (**B**) Psychometric curves for marmosets H and I (mean ± SEM). (**C**) Temporal weighting curves for both marmosets in the DSTA task. The horizontal axis is the temporal position of each tone, and the vertical axis is the psychophysical weight of each tone, which represents the contribution to the final decision.

Psychometric curves showed that all marmosets successfully learned the DSTA task (Figure 2B). Moreover, stimuli with the same evidence strength but different sequence patterns exhibited distinct choice accuracy (Supplemental Figure 1A). Both marmosets maintained a low violation rate in the task (Supplemental Figure 1C).

The reverse correlation analysis showed that early tones contributed more to decision-making in this complex decision-making task (Figure 2C).

### Accumulation strategy dominates marmoset decision-making

In the DSTA task, besides integrating multiple tones sequentially, subjects could also adopt a simpler strategy: making decisions based on a single, randomly selected tone. This single-sample strategy does not require integration, but can still yield above-chance performance, especially for static-evidence stimuli, in which all tones point toward the correct choice. In such cases, if the internal signal-to-noise ratio (SNR) for single tone discrimination is sufficiently high, it becomes difficult to distinguish this strategy from multi-sample integration. However, for dynamic-evidence stimuli, in which tone frequencies signaling correct or erroneous choice randomly alternate throughout the stimulus, even with a high internal SNR for single tone discrimination, the single-sample strategy yields markedly lower performance relative to multi-sample integration.

Based on the above theoretical discussion, we used computational modeling to disentangle the integration strategy from the single-sample strategy(Hyafil et al., 2023; Stine et al., 2020). For the integration strategy, we developed an integration model, in which the final decision was based on the weighted integration of multiple tones (see Methods, Eqs. 1–2). For the single-sample strategy, we developed a snapshot model in which the final choice was determined by sum of the selection probabilities of tones with the same frequency in the stimulus. (see Methods, Eqs. 3–6). We separately fitted the integration model and snapshot model to the marmoset behavioral data and compared the task performance between the model and the marmoset behavior. We found that integration model is better fit to the marmoset behavior (Figure 3A). For the integration model, the psychometric and temporal weighting curves nearly overlapped with those of marmoset behavior (Figure 3B, C). For the snapshot model, performance on static-evidence stimuli matched marmoset behavior well, whereas performance on dynamic-evidence stimuli fit poorly (Figure 3D), which is consistent with the theoretical analysis above. Moreover, although the temporal weighting curves of the snapshot model and marmosets were similar in shape, the model’s weights at each tone position were systematically lower than those of marmosets (Figure 3E). Collectively, these results indicate that the marmosets indeed accumulated multiple pieces of evidence for decision-making.

**Figure 3.**
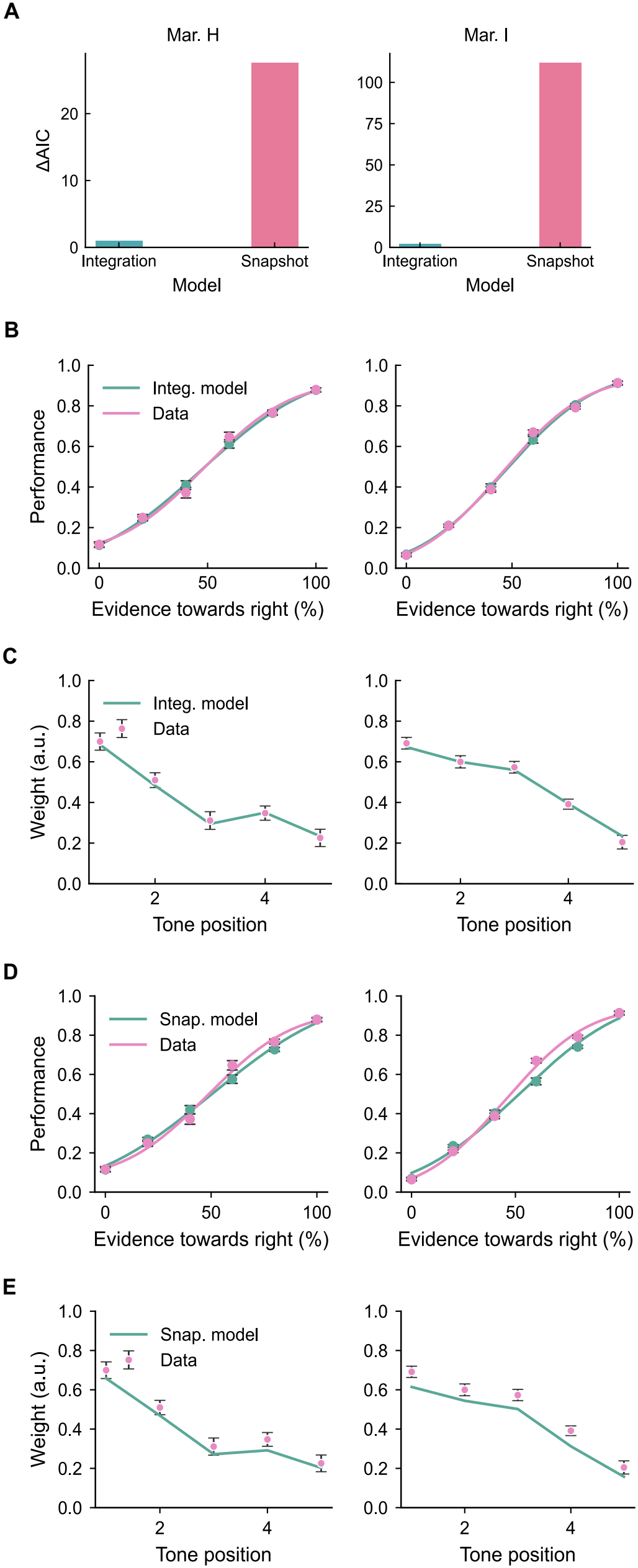
Integration model and snapshot model comparison results. (**A**) Model comparison via Akaike Information Criterion (AIC). The horizontal axis represents model types, the vertical axis shows the difference in AIC values relative to the integration model (baseline), with higher values indicating worse model fit. (**B, C**) Psychometric and temporal weighting curves for marmoset behavior (pink) and integration model (green). (**D, E**) Psychometric and temporal weighting curves for marmoset behavior (pink) and snapshot model (green). (**A**) Comparison of temporal weighting curves for 80% (green) and 60% (orange) evidence strength (Wald test). Evidence strength was defined by the proportion of tones belonging to the majority category (e.g., 80% = 4 tones of one frequency and 1 tone of the other). Error bars represent 95% confidence intervals. †: *P* < 0.1, *: *P* <0.05, **: *P* < 0.01, ***: *P* < 0.001. For marmoset H, the *P* value for each tone position is: *P*_1_ = 0.118, *P*_2_ = 0.689, *P*_3_ = 0.0498, *P*_4_ = 0.0005, *P*_5_= 0.0514; For marmoset I, the *P* value for each tone position is: *P*_1_ = 0.598, *P*_2_ = 0.377, *P*_3_ = 0.0508, *P*_4_ = 0.00320, *P*_5_= 0.106. The subscript numbers denote tone positions.

### Psychophysical weights reveal the ETW effect

Bounded evidence accumulation mechanism predicts that lower evidence strength leading to the contribution of late evidence increase and thus producing the ETW effect. To test this prediction, we used logistic regression model with interaction terms(Hosmer et al., 2013) to analyze the psychophysical weights for 80% and 60% evidence strength (see Methods, Eqs. 7). Consistent with the prediction, we found the weights for late tones were higher under 60% evidence strength than under 80% evidence strength in both marmosets (Figure 4A). This analysis quantitatively revealed the ETW effect in the DSTA task.

Besides this fixed-duration version of the DSTA task, we also developed a reaction-time version of the DSTA task in marmosets, in which the marmosets were free to respond after stimulus onset (Supplemental Figure 2A). The marmosets performed the reaction-time task well (Supplemental Figure 2B, C). Furthermore, the reaction-time version also showed the ETW effect (Supplemental Figure 2D), albeit weaker than that observed in the fixed-duration version.

**Figure 4.**
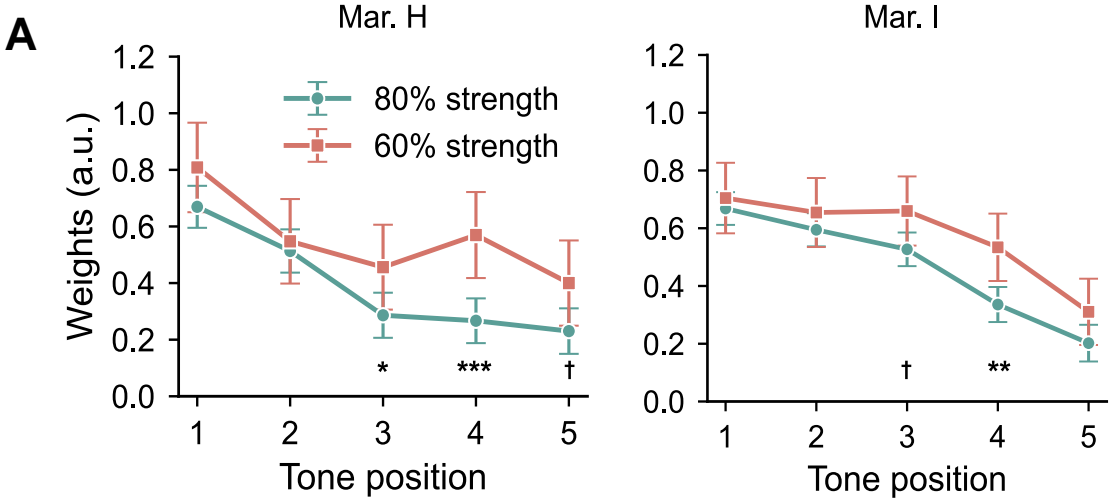
Temporal weighting curves for different evidence strengths. **(A)** Comparison of temporal weighting curves for 80% (green) and 60% (orange) evidence strength (Wald test). Evidence strength was defined by the proportion of tones belonging to the majority category (e.g., 80% = 4 tones of one frequency and 1 tone of the other). Error bars represent 95% confidence intervals. †: *P* < 0.1, *: *P* <0.05, **: *P* < 0.01, ***: *P* < 0.001. For marmoset H, the *P* value for each tone position is: *P*_1_ = 0.118, *P*_2_ = 0.689, *P*_3_ = 0.0498, *P*_4_ = 0.0005, *P*_5_= 0.0514; For marmoset I, the *P* value for each tone position is: *P*_1_ = 0.598, *P*_2_ = 0.377, *P*_3_ = 0.0508, *P*_4_ = 0.00320, *P*_5_= 0.106. The subscript numbers denote tone positions.

### Candidate behavioral mechanisms to account for the ETW effect

The standard drift diffusion model is a canonical bounded accumulation model. In this model, it is assumed that the accumulator equally accumulates momentary evidence under constant diffusion noise until the accumulated evidence reaches a decision bound. Following this bounded accumulation framework, the observed ETW effect in the DSTA task should be attributed to the greater incorporation of late tones into the decision when evidence strength is weaker. Therefore, we developed a DDM with constant diffusion noise and identical gain modulation for each accumulated tone (CDN-IG model) to test that whether bounded accumulation mechanism alone is sufficient to account for the observed ETW effect (see Methods). Surprisingly, the best fitting model showed that the temporal weighting curves for 80% and 60% evidence strength almost overlapped (Figure 5A), indicating the bounded accumulation mechanism alone was insufficient to account for the ETW effect.

**Figure 5.**
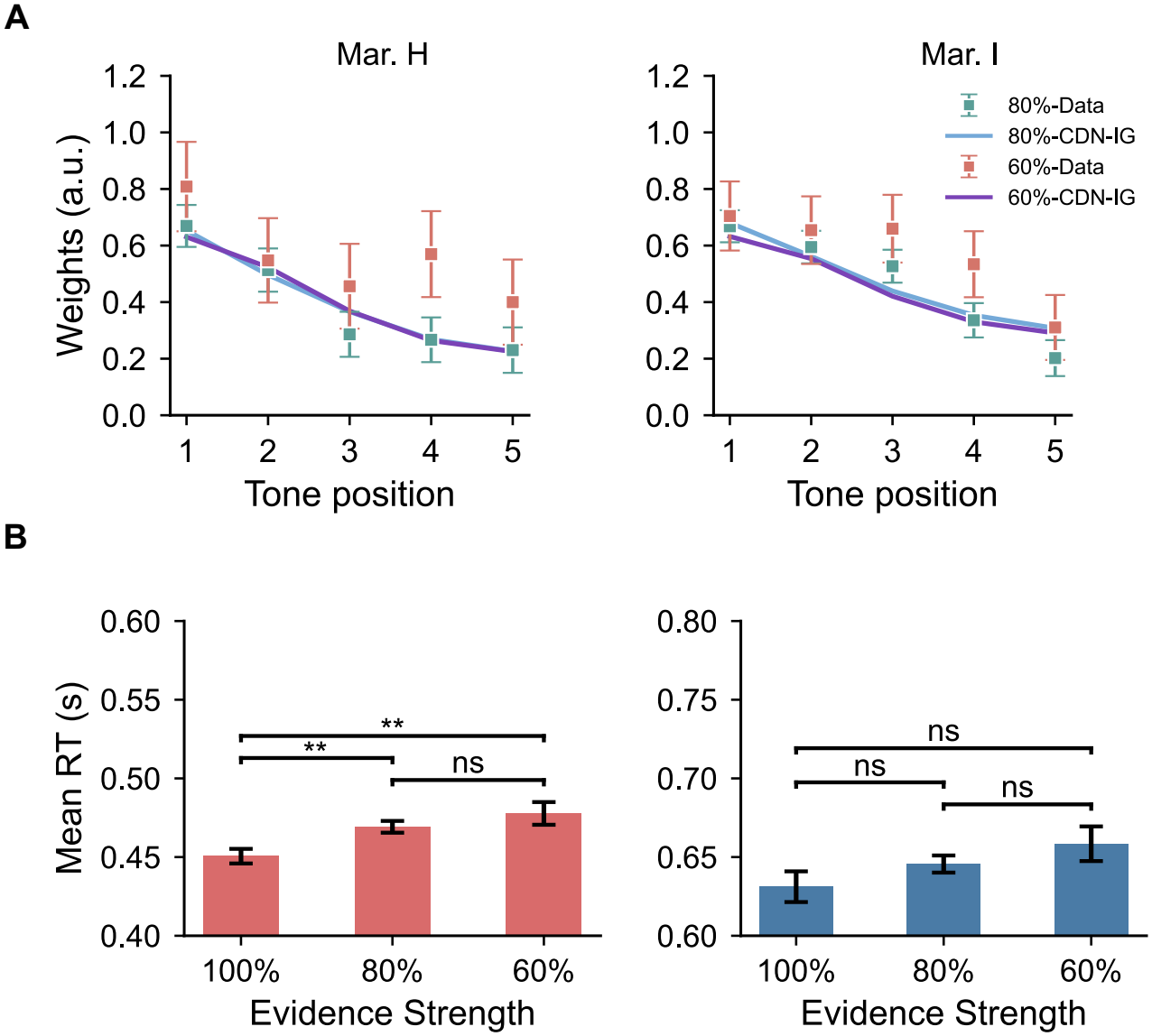
Candidate behavioral mechanisms to account for the ETW effect. (**A**) Curves show the psychophysical weights from the CDN-IG model for 80% (Data: green dots; Model: blue curve) and 60% (Data: orange dots; Model: purple curve) evidence strength conditions. The Wald test was used to compare weights between the two conditions in CDN-IG model. For marmoset H, the *P* value for each tone position is: *P*_1_ = 0.795, *P*_2_ = 0.755, *P*_3_ = 0.979, *P*_4_ = 0.961, *P*_5_= 0.992; For marmoset I, the *P* value for each tone position is: *P*_1_ = 0.478, *P*_2_ = 0.902, *P*_3_ = 0.779, *P*_4_ = 0.725, *P*_5_= 0.801. All tone positions are not significant. Dots with error bars show the observed psychophysical weights from marmoset behavioral data. All error bars denote 95% confidence intervals. (**B**) Bars represent mean RT after go cue at each evidence strength level (100%, 80%, 60%); error bars denote the standard error of the mean (SEM). Pairwise comparisons were performed using Welch’s independent samples t-tests, with Bonferroni correction applied to control for multiple comparisons (corrected significance threshold α = 0.05/3 ≈ 0.017). Significance markers: * *p* < 0.017, ** *p* < 0.01, *** *p* < 0.001; ns, not significant. For marmoset H, 100% vs 80%: *P* = 0.0018, 100% vs 60%: *P* = 0.0016, 80% vs 60%: *P* = 0.295; For marmoset I, 100% vs 80%: *P* = 0.197, 100% vs 60%: *P* = 0.0636, 80% vs 60%: *P* = 0.294.

To explain the observed ETW effect in the DSTA task, we further proposed the mechanism of sampling from memory(Shushruth et al., 2022). For weaker evidence strength, there are more trials that the DV has not reached the decision bound by the end of the stimulus. In such a case, the brain may continue to sample and accumulate evidence from memory. With memory decay, early evidence is more likely to be forgotten, causing late evidence to be sampled more frequently. This in turn increases the contribution of late evidence to decision-making, producing the ETW effect. In the DSTA task, the go cue was presented 0.1 s after stimulus offset. If the marmosets sampled from memory more frequently under weaker evidence strength, the response time after the go cue should be significantly longer in the 60% evidence-strength condition than in the 80% condition. However, our analysis revealed that although response times showed a slight increasing trend, there was no significant difference between the two conditions (Figure 5B), suggesting that marmosets did not rely on post-stimulus memory sampling to generate the ETW effect in the DSTA task.

### DDM variant with evidence-strength-dependent diffusion noise successfully reproduce the ETW effect

In the DSTA task, sensory inputs and motor outputs are tightly controlled, which greatly constrains the space of possible mechanisms underlying the ETW effect. After above candidate mechanisms are excluded, the ETW effect are most likely to be attributed to the adaptive modulation of cognitive state as perceived task difficulty increases. The cognitive state can be decision-making caution or cognitive effort (e.g., attentional engagement with sensory evidence). In the bounded accumulation framework, greater caution implies a higher decision bound, leading to more late tones being incorporated into the decision and thus higher weights for late tones. The logic underlying this mechanism is similar to that in the standard DDM described above. We built a DDM variant with evidence-strength-dependent decision bounds. As in the CDN-IG model, this DDM variant also failed to account for the ETW effect (Supplemental Figure 3A).

Therefore, the adaptive modulation of cognitive effort constitutes the most viable mechanism underlying the ETW effect. This interpretation is reasonable when considering the actual task context. For difficult stimuli, highly motivated marmosets are eager to obtain rewards quickly while sustaining choice accuracy. Post-stimulus memory sampling would delay the reward acquisition, conflicting with internal urgency. In such a case, a natural solution is to increase cognitive effort during decision-making, thereby suppressing neural noise or amplifying sensory evidence via gain modulation to sustain task performance. For weaker evidence strength, as the tone sequence unfolds, the more frequent alternation between low and high frequency tones causes the perceived task difficulty to gradually increase relative to stronger evidence strength. The marmosets therefore adaptively enhance their attentional engagement with late tones, resulting in higher weights for late tones.

In the bounded accumulation framework, the modulation of cognitive effort across evidence strengths may be modeled as evidence-strength-dependent diffusion noise or evidence-strength-dependent gain modulation. The evidence-strength-dependent gain modulation will globally modulate the contribution of both signal and noise tones to the decision since it is unknown which tone frequency is the signal during evidence accumulation, and thus it is unable to increase the weight of late tones for weaker evidence strength during decision-making.

The best-fitting DDM variant with evidence-strength-dependent gain modulation also failed to capture the ETW effect (Supplemental Figure 3B). Therefore, we set the diffusion noise to be evidence-strength-dependent while keeping the gain modulation constant across evidence strengths. In addition, to capture the observed non-monotonic temporal weighting patterns, we introduced a tone-position-dependent gain modulation mechanism into the model. In this way, we constructed an DDM variant with evidence-strength-dependent diffusion noise (EDN model, see Methods, Eqs. 9-10) to capture the observed ETW effect in the DSTA task. During model fitting, we fixed the diffusion noise for the 100% evidence strength at a constant value of 5. Under this constraint, the gain modulation parameters could be identified. Because the gain modulation parameters were constant across evidence strengths, the remaining diffusion noise parameters were also identifiable.

The EDN model produced robust fits to the behavioral data, and the estimated parameters were highly consistent across multiple fits from random initializations (Supplemental Figure 4). The best-fitting model precisely captured the psychometric and temporal weighting curves (Supplemental Figure 5). Notably, it closely matched the observed choice accuracy across all sequence patterns in the behavioral data (Figure 6A). Further psychophysical weight analysis (as in Figure 4) also revealed greater weights of late tones for lower evidence strength (Figure 6B), indicating that the model successfully reproduced the ETW effect in the DSTA task.

**Figure 6.**
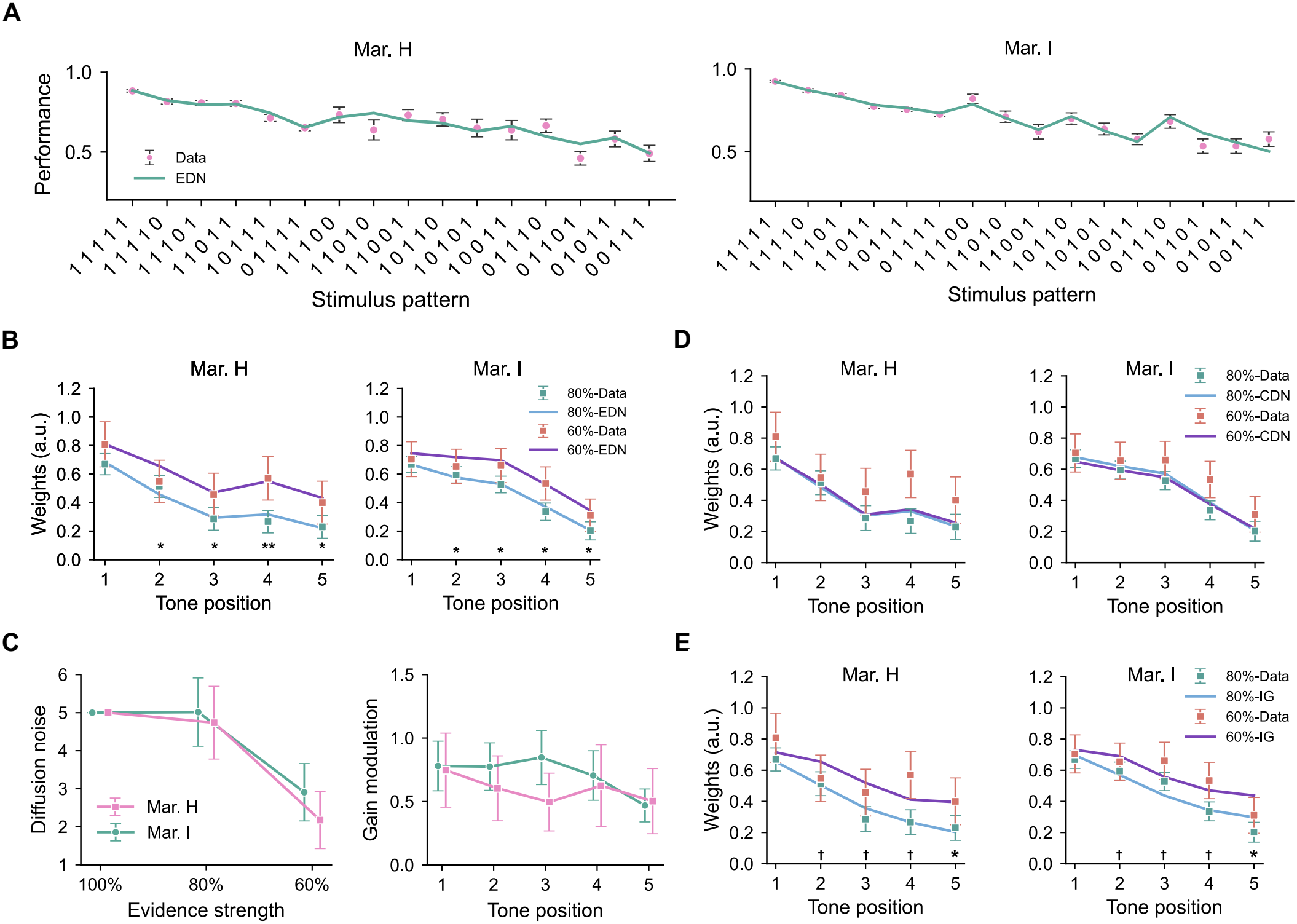
Evidence-strength-dependent diffusion noise is a key computational component to reproduce the ETW effect. (**A**) Choice accuracy for each sequence pattern from marmoset behavior (pink) and from the EDN model (green). The horizontal axis represents the sequence patterns, where 1 and 0 denote signal (correct) and noise (erroneous) tones, respectively. (**B**) Psychophysical weights from the EDN model and marmoset behavior for the 80% (Data: green dots; Model: blue curve) and 60% (Data: orange dots; Model: purple curve) evidence-strength conditions. Error bars represent 95% confidence intervals, and asterisks indicate significant differences between the two conditions in the model (Wald test). For marmoset H, the *P* value for each tone position is: *P*_1_ = 0.159, *P*_2_ = 0.0222, *P*_3_ = 0.0422, *P*_4_ = 0.00761, *P*_5_= 0.0157; For marmoset I, the *P* value for each tone position is: *P*_1_ = 0.254, *P*_2_ = 0.0364, *P*_3_ = 0.0146, *P*_4_ = 0.0188, *P*_5_= 0.0400. (**C**) Estimated values of diffusion noise parameters for each evidence strength (left) and gain modulation parameters for each tone position (right). Error bars represent standard errors derived from the inverse Hessian matrix. The diffusion noise for 100% evidence strength was fixed to 5 during model fitting. (**D**) The same as in panel (**B**), but for CDN model. The CDN model is a DDM with constant diffusion noise and tone-position-dependent gain modulation. For marmoset H, the *P* value for each tone position is: *P*_1_ = 0.555, *P*_2_ = 0.844, *P*_3_ = 0.657, *P*_4_ = 0.894, *P*_5_= 0.823; For marmoset I, the *P* value for each tone position is: *P*_1_ = 0.649, *P*_2_ = 0.750, *P*_3_ = 0.706, *P*_4_ = 0.928, *P*_5_= 0.771. All tone positions are not significant. (**E**) The same as in panel (**B**), but for the IG model. The IG model is a DDM with evidence-strength-dependent diffusion noise and identical gain modulation across tone positions. The daggers and asterisks represent the results of comparisons between the weights for the 80% and those for the 60% evidence-strength conditions in the IG model. For marmoset H, the *P* value for each tone position is: *P*_1_ = 0.496, *P*_2_ = 0.0804, *P*_3_ = 0.0597, *P*_4_ = 0.0899, *P*_5_= 0.0276; For marmoset I, the *P* value for each tone position is: *P*_1_ = 0.611 *P*_2_ = 0.0761, *P*_3_ = 0.0742, *P*_4_ = 0.0560, *P*_5_= 0.0368 (Wald test).

The best-fitting model further revealed that the diffusion noise decreased for lower evidence strengths, while the gain modulation fluctuated across tone positions (Figure 6C). In the bounded accumulation framework, the lower diffusion noise implies that DV evolution is subject to weaker random perturbations, leading to higher choice accuracy. Under the constraint of behavioral data, the EDN model naturally yielded lower diffusion noise to sustain choice accuracy for weaker evidence strengths, which was consistent with the adaptive modulation of cognitive effort mechanism.

By constraining the diffusion noise to be constant across evidence strengths while allowing the gain modulation to fluctuate across tone positions, we built a constant diffusion noise (CDN) model. This model fit was worse than that of the EDN model and it failed to capture the ETW effect (Figure 6D). Conversely, by constraining the gain modulation to be identical across tone positions while allowing the diffusion noise to vary with evidence strengths, we built the identical gain (IG) model. This model still captured the ETW effect (Figure 6E). Since the gain modulation was identical across tone positions in this model, the temporal weighting curve was monotonic and thus failed to capture the non-monotonic temporal weighting patterns. These modeling results further indicate that the evidence-strength-dependent diffusion noise is the key computational component for generating the ETW effect in the EDN model.

## Discussion

### Bridging the gap between higher cognition and circuit-level investigation: cognitive task training in the marmoset

Elucidating how projection patterns and neural dynamics govern higher cognition requires mechanistic investigation at the neural circuit level. While rhesus monkeys have served as the primary non-human primate model in cognitive neuroscience, neural circuit-level investigation has been limited, because their large body size, deeply folded cortex, and high experimental costs make it difficult to apply advanced circuit analysis techniques(Cushnie et al., 2020; Galvan et al., 2017). Although these circuit tools have been intensively used in rodents, rodents lack the cognitive complexity of primates. As a small non-human primate species, marmosets offer a promising alternative to bridge this gap. However, the absence of complex cognitive behavioral paradigms for head-restrained marmosets has long constrained their use in cognitive neuroscience(Okano, 2021). To our knowledge, this study is the first to show that head-restrained marmosets can perform complex cognitive tasks comparable to those well-established for macaques, indicative of robust cognitive capacities in marmosets. Given the rapid development of neural circuit analysis techniques for marmosets(Dotson et al., 2024; Jendritza et al., 2023; Kondo et al., 2018; Majka et al., 2020; Shaw et al., 2026; Walker et al., 2021), this work provides a key foundation for unraveling neural circuit mechanisms underlying higher cognition in primates.

### Adaptive modulation of cognitive effort in complex decision-making

In the standard DDM, the ETW effect originates from the extended accumulation period for weaker evidence strength, which provides greater opportunity for late evidence to be integrated into DV(Kiani et al., 2008; Ratcliff and McKoon, 2008). This explanation assumes that the drift rate is determined by the perceptual sensitivity to the stimulus, while the diffusion noise is held constant(Palmer et al., 2005; Ratcliff et al., 2016). The perceptual sensitivity to the stimulus may be modulated by attention(Moore and Zirnsak, 2017; Reynolds and Chelazzi, 2004), but this mechanism cannot be identified at the behavioral level in many decision-making paradigms (such as random dot kinematogram task). In the DSTA task, since the tone intensity is identical across all evidence strengths, if there is no sensory adaptation or cognitive modulation to the perceptual sensitivity, the perceptual sensitivity should be identical for the same tone frequency across all evidence strengths. As a feedforward mechanism at the sensory level, sensory adaptation cannot explain the ETW effect(Kawaguchi et al., 2018). Therefore, in the standard DDM, the observed ETW effect in our task can only be attributed to the greater incorporation of late tones into the decision for difficult stimuli. This explanation is intuitively plausible. However, our modeling results show that only the bounded accumulation mechanism is insufficient to generate the ETW effect in the DSTA task. This finding may stem from the stimulus design. For 80% evidence strength, there is one noise tone in the tone sequence, which can appear at any position. If marmosets consistently rely on early tones to make decisions and ignore late tones, stimuli with noise tone in the early position will mislead the animals. Therefore, these stimuli drive marmosets to incorporate late tones into the decision more frequently. For 60% evidence strength, many stimuli require using all 5 tones to achieve correct choices. Given the limited working memory capacity(Zlatkina et al., 2024), it is cognitively challenging for marmosets to use all 5 tones to make decisions. As a result, late tones are used less frequently, even though they should be used more in the 60% evidence strength. These two factors jointly make the number of tones incorporated into decisions similar across the two evidence-strength conditions, which further causes the bounded accumulation mechanism alone to fail to account for the ETW effect.

The DSTA task tightly constrains the space of possible behavioral strategies through its quantitatively controlled sensory inputs and motor outputs. Within this highly constrained task environment, our statistical and model-based analyses collectively reveal that the ETW effect arises primarily from adaptive modulation of cognitive effort (such as the attentional engagement with the stimulus) during evidence accumulation. In the bounded accumulation framework, this cognitive modulation manifests as evidence-strength-dependent diffusion noise. This adaptive modulation may stem from the real-time monitoring of task difficulty during decision-making. The metacognitive monitoring drives the brain to actively suppress internal neural variability for difficult trials, thereby improving the signal-to-noise ratio for each accumulated sample and enabling more precise evidence accumulation. Recent studies in humans and macaques have shown the online confidence monitoring during decision-making(Balsdon et al., 2020; Dotan et al., 2018; Vivar-Lazo and Fetsch, 2026), raising the possibility that metacognitive monitoring drives the adaptive modulation of attentional engagement with the sensory evidence. A large body of work has established that attentional modulation directly shapes internal neural variability(Cohen and Maunsell, 2009; Mitchell et al., 2009; Moore and Zirnsak, 2017). Future work should employ advanced neural circuit analysis techniques to test these predictions and further elucidate the neural computational mechanisms underlying discrete evidence accumulation in primates.

## Methods

### Animals

Four adult common marmosets (*Callithrix jacchus*) between the ages of 2–5 years were used in this study. All experiments were approved by the Animal Ethics Committee of the Center for Excellence in Brain Science and Intelligence Technology, Chinese Academy of Sciences (ION-2019019), and Animal Welfare and Use Committees of the Shenzhen Institutes of Advanced Technology, Chinese Academy of Sciences (IAT-BSI-IRB-NHP-202401030-WUXXL-A0028). During task training, marmosets lay prone in a body-restraint tube. Marmoset B was implanted with a customized head plate for head fixation; marmosets H, I, and K did not undergo surgery, and their heads were restrained by customized helmets. Animals were housed under a 12-h light/dark cycle and fed a commercial diet (Jiangsu Xietong Pharmaceutical Bio-engineering Co., Ltd.) accompanied by fruits, cookies, and marshmallows. Water intake was restricted before training to motivate task performance. Body weight was monitored daily and maintained within 85–95% of the average weight before water restriction.

### Surgery

Marmoset B fasted for 12 hours before surgery. The animal was then anesthetized with 1–2% isoflurane (RWD Life Science). Atropine (0.1 mg/kg), ampicillin (40 mg/kg), and dexamethasone (0.5 mg/kg) were administered intramuscularly to reduce secretions and prevent inflammation before surgery. Then, the marmoset’s head was fixed on a stereotaxic frame (Narishige). Before incising the scalp, 0.4% lidocaine was applied for local analgesia at the intended incision site. A customized titanium head plate and skull screws were implanted on the skull. During surgery, the animal’s heart rate, blood oxygen saturation, respiratory rate, and body temperature were monitored using a medical monitor (Mindray). The heart rate was generally maintained between 170–200 beats per minute (bpm), and body temperature was kept at approximately 37°C using a heated blanket(Mundinano et al., 2016). After surgery, meloxicam (0.2 mg/kg) was also injected intramuscularly for postoperative analgesia.

### Behavioral setup

The behavioral setup was controlled by a development board (Arduino Mega 2560 or Arduino GIGA R1 WiFi). Auditory stimuli were amplified and delivered via a commercial power amplifier and speakers (Tucker-Davis Technologies, TDT or PAIYON Audio Co., Ltd.)(Zhong et al., 2019). Behavioral tasks were controlled by custom scripts executed on the Arduino microcontroller. The marmoset restraint device and the soundproof chamber were custom-built. The restraint device consisted of a 3D-printed body restraint tube, a head-restrained plate, and a helmet.

### Behavioral tasks

#### PFD and DGC tasks (Marmoset B)

Marmoset B was trained on the pure tone frequency discrimination (PFD) task and the delayed go-cue (DGC) task, respectively. In the PFD task, each trial began with illumination of a white LED to signal task availability. The marmoset licked the central spout to initiate the trial, after which the LED was turned off. Following a random delay (0.5–1 s), a 0.5 s pure tone was presented at 65 dB. A high-frequency tone (12.7 kHz) signaled a rightward choice; a low-frequency tone (4.5 kHz) signaled a leftward choice.

Training proceeded in four stages. (1) Immediate reward: Licking either the left or right spout delivered 10–20 μL of water. After 20–40 licks on one side, the marmoset was required to switch to the other spout to prevent side biases. (2) Side-matched tone: Licking a spout triggered a tone matching that side; reward was delivered only after tone offset. The required side alternated every 3–5 trials, and reward volume was higher than in the first stage to encourage licking. (3) Frequency discrimination: High and low tones were presented randomly across different trials; the marmoset learned the frequency–choice mapping through trial and error over 2–4 weeks. (4) Trial initiation: A central spout was added. Marmosets first learned to lick the central spout to obtain reward from the side spouts. Subsequently, a white LED was turned on at trial onset. If the marmoset licked the central spout while the LED was on, the LED was turned off and a reward was delivered. Licking while the LED was off had no effect. Once this rule was mastered, the tone discrimination task was reintroduced. In this protocol, tone discrimination was learned before trial initiation; reversing the order was also effective.

The DGC task was similar to the PFD task, except that a random delay (0.5–0.8 s) was inserted between stimulus offset and the response period. A red LED signaled the end of the delay and permitted a response. Premature licks during this random delay extended the waiting period until licking ceased.

#### Fixed-duration (FD) version of the DSTA task (Marmosets H and I)

Marmoset H and I were trained on the dynamic sequential tone accumulation (DSTA) task. The DSTA task resembled the DGC task but used a sequence of five pure tones, each randomly assigned a low (5.4 kHz) or high (11.8 kHz) frequency. Each tone lasted 0.125 s, with a 0.125 s inter-tone interval, yielding a total stimulus duration of 1.25 s. Marmosets were required to report which frequency predominated in the tone sequence. During stimulus presentation, marmosets had to withhold responses until stimulus offset and a subsequent go cue. The interval between stimulus offset and go cue onset was 0.1 s; the go cue was a 0.15 s exponential frequency sweep ranged from 5.4 kHz to 11.8 kHz. The proportion of difficult trials was adjusted to an appropriate level to control overall task difficulty (Supplemental Figure 1B). Licking before the go cue resulted in two penalties: the delay period between stimulus offset and go cue was prolonged, and the reward volume was reduced.

#### Reaction-time (RT) version of the DSTA task (Marmosets I and K)

Marmosets I and K were trained on a reaction-time version of the DSTA task. In this task version, subjects were free to report their decisions after the stimulus onset. To balance the speed-accuracy trade-off, as in prior studies(Kira et al., 2015; Stine et al., 2023), a reward delay period (1.2 s) was imposed after stimulus onset. Correct responses in this period were rewarded only after the delay elapsed, while correct responses after this period led to immediate rewards.

### Behavioral data preprocessing

For the fixed-duration task version, only sessions with performance on non-violation trials above approximately 75% were included for data analysis and modeling. Only non-violation trials from these included sessions were used for analysis, while violation trials and miss trials were excluded. Based on this criterion, a total of 20020 trials were collected, with marmoset H contributing 7,626 trials and marmoset I contributing 12,394 trials. For the reaction-time task version, only sessions with performance above 70% were included, and miss trials were excluded. This criterion yielded a total of 19,695 trials, with marmoset I contributing 8,016 trials and marmoset K contributing 11,679 trials. All behavioral data analysis and modeling were performed using customized Python

(Python 3.12) scripts.

### Integration model optimization and simulation

The integration model was an extended logistic regression model(Hyafil et al., 2023). For the integration strategy, we developed an integration model, in which the final decision was based on the weighted integration of multiple tones:

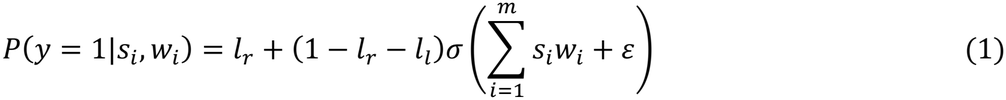

In this equation, *i* indexes the *i*^*th*^ tone sample, and *m* denotes the total number of tone samples within a single stimulus. *l*_*r*_ and *l*_*l*_ represent the rightward and leftward lapse rates, respectively. *w*_*i*_ is the psychophysical weight assigned to the *i*^*th*^ tone position. *s*_*i*_ denotes the sample value; *s*_*i*_ = ™1 corresponds to a low frequency, and *s*_*i*_ = 1 corresponds to a high frequency. *ε* denotes the bias term, and *σ* denotes the logistic function.

Maximum likelihood estimation (MLE) was adopted to estimate the weight parameter *w*_*i*_, which quantified the psychophysical weight of each tone position during evidence integration. The log-likelihood function was defined as:

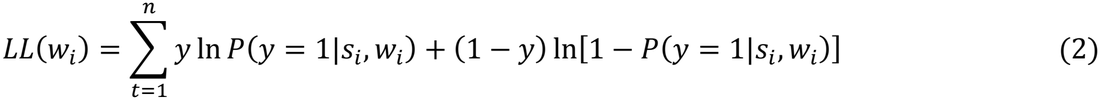

During model fitting, the lapse parameters *l*_*r*_ and *l*_*l*_ were directly derived from the error rates of stimuli with 100% evidence strength (all-high-frequency and all-low-frequency stimuli) in behavioral data. Hence, these two parameters were fixed during model fitting. Since the lapse rate was independent of sensory evidence, sensory evidence strength was assigned negative infinity for all-low-frequency stimuli and positive infinity for all-high-frequency stimuli during model fitting. The term *P*(*y* = 1|*s*_*i*_, *w*_*!*_) was calculated according to equation (4). After model fitting and estimation of individual *w*_*i*_ values, behavioral simulations were performed using the optimized parameters. To obtain stable simulation results, behavioral simulations were repeated 100 times using the same stimulus set as in the marmoset behavioral task.

### Snapshot model optimization and simulation

For the snapshot model, a single sample in each tone sequence was randomly selected to guide decision-making. If the sampled evidence indicated a correct choice, the marmosets would respond correctly, and vice versa. The snapshot model is given by:

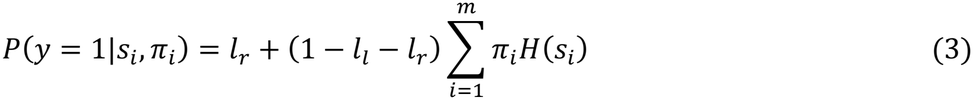

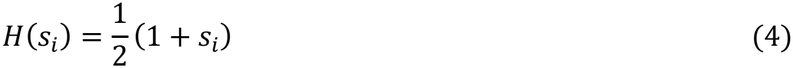

The function *H*(*s*_*i*_) maps each tone sample onto the corresponding choice. *π*_*i*_ denotes the selection probability of the *i*^*th*^ tone within the stimulus, with the sum of *π*_*i*_ equal to 1. Similar to the integration model, the lapse rate in the snapshot model was defined according to the error rate of stimuli with 100% evidence strength. The parameter *π*_*i*_ was estimated via constrained optimization. The constrained optimization is formulated as:

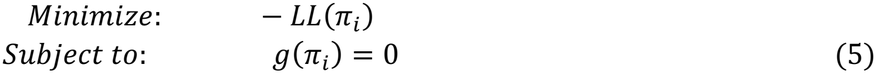

The *LL*(*π*_*i*_) is defined similarly to equation (5), while *g*(*π*_*i*_) is the constraint condition.

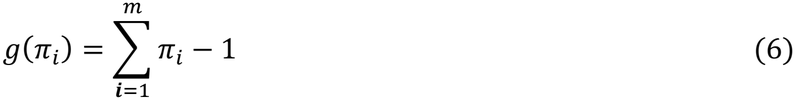

Sequential Least Squares Programming (SLSQP, an optimization algorithm available in the Python Scipy package) was used for model optimization and *π*_*i*_ estimation. During model fitting, the lapse parameters *l*_*r*_ and *l*_*l*_ were fixed, consistent with the integration model. After estimating the *π*_*i*_ values, 100 independent behavioral simulations were performed using the snapshot model to obtain stable comparison results between the model and marmoset behavior. The comparison approach was the same as that used in the previous integration model.

### Psychophysical weights analysis

Psychophysical weights for each tone position were estimated via logistic regression. The logistic regression was fitted via the Logit class implemented in the statsmodels package. For each marmoset behavioral data set, a logistic regression model was fitted to behavioral data from individual training sessions to extract session-wise psychophysical weights. Final weights were then averaged across all sessions. The logistic regression model with interaction terms was used to analyze the psychophysical weights under different evidence strengths. This model was formulated as

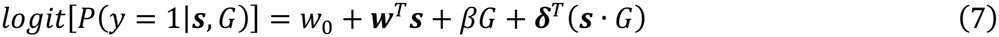

Where *s* = (*s*_1_, *⋯* , *s*_*m*_)^*T*^ denoting the tone sequence, in which *s*_*i*_ takes a value of 1 (for high-frequency tones) or -1 (for low-frequency tones), and *m* is the total tone number in the sequence. *G* is a dummy variable denoting the evidence strength ( *G* = 0 for the 80% evidence strength; *G* = 1 for the 60% evidence strength). *w*_0_ is the intercept for 60% evidence strength, and the vector *w* = (*w*_1_, *⋯* , *w*_*m*_)^*T*^ representing the psychophysical weights for the 5 tones in the 60% evidence strength. The *β* captures the difference in intercept between 80% and 60% evidence strength. The interaction coefficients *δ* = (*δ*_1_, *⋯* , *δ*_*m*_)^*T*^ quantify how the weights of the tones differ between the two evidence strengths. The weight of tone *s*_*i*_ in 60% evidence strength is *w*_*i*_ + *δ*_*i*_. Consequently, *δ*_*i*_ directly measures the change in the weight assigned to tone *s*_*i*_ when moving from 80% to 60% evidence strength.

### Drift diffusion models (DDM) optimization

The temporal evolution of DV in the DDM with evidence-strength-dependent diffusion noise (EDN model) is defined in equations (8) and (9). Based on this, the CDN model constrained the diffusion noise to be a constant value of 5, the IG model constrained the gain modulation to be identical across tone positions (gain modulation value was set to 1), and the CDN-IG model imposed both constraints (The diffusion noise parameter was modeled as a free parameter, and the gain modulation was set to 1). The DDM with evidence-strength-dependent decision bounds (EDB model) was constructed by constraining the diffusion noise to a constant value of 5 across different evidence strengths and by separately parameterizing the decision bound for each evidence strength. Tone-position-dependent gain modulation was also incorporated into this DDM variant. The DDM with evidence-strength-dependent gain modulation (EGM model) was constructed by setting the diffusion noise as a free parameter and by separately parameterizing the gain modulation for each evidence strength. For model identifiability, the gain modulation for 100% evidence strength is set to 1. The gain modulation across different tone positions is set to identical in this DDM variant.

In this EDN model, when the DV is in the decision bound, the temporal evolution of the DV is modeled as

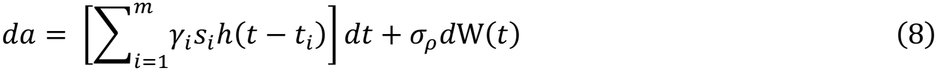

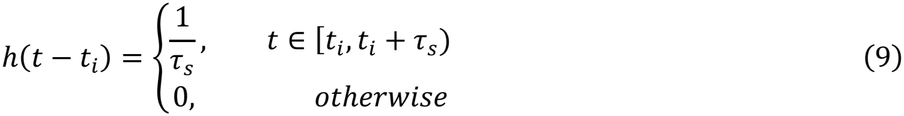

Here, *a* denotes the DV. *γ*_*i*_ represents the gain modulation of the *i*^*th*^ accumulated tone to the DV increment. *h*(*t* ™ *t*_*i*_) is a normalized uniform kernel function, where *τ*_s_ is the duration of each tone and *t*_*i*_ denotes the onset time of the *i*^*th*^ tone. This function distributes each discrete tone evenly over its duration and enables discrete evidence to align with the continuous evidence accumulation process of the DDM. *σ*_*ρ*_ represents the diffusion noise, and *ρ* denotes the evidence strength.

Model optimization was adapted from the DDM described in Brunton et al. Maximum likelihood estimation (MLE) was adopted for model fitting, with the log-likelihood function expressed as:

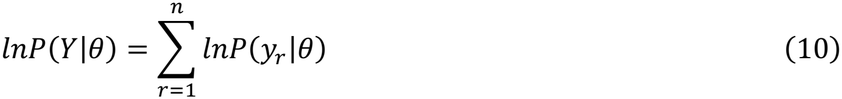

Here, *n* indicates the total number of trials; *r* denotes the trial index; *θ* represents the full parameter set; and *Y* represents the choice set. The expression of *lnP*(*y*|*θ*) is required to construct the log-likelihood function. Since choice *y* is decided by the DV at the stimulus offset time point *T*, the choice probability is defined as:

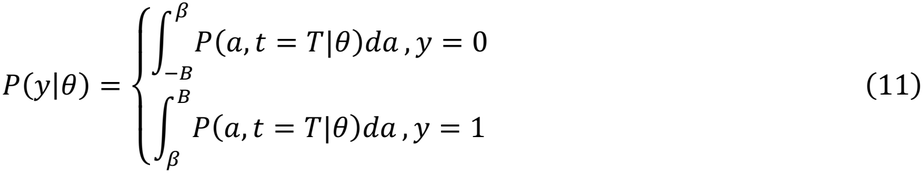

Here, *P*(*a, t* = *T*|*θ*) is the DV probability distribution at the final time step. *β* denotes the bias, and *B* denotes the decision bound. Both time t and DV value *a* are discretized, with *a* = {*ξ*_1_, *ξ*_2_, … , *ξ*_p_, … *ξ*_*N*_}. The deterministic component in equation (4) is:

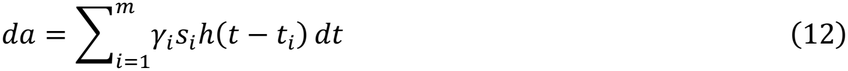

After discretization, the average DV value at time step *k*, derived from the DV value at previous time step *k* ™ 1, is given by:

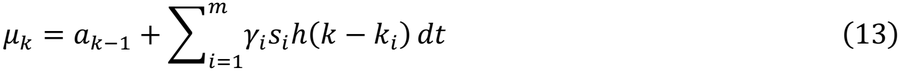

where:

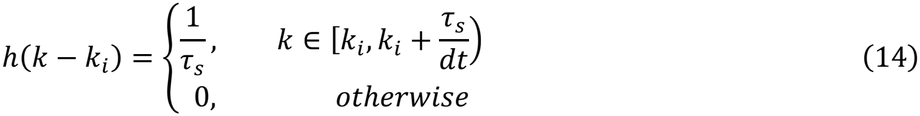

During DV evolution, the transition probability from *a*_*k−1*_ *t*o *a*_*k*_ can be represented as:

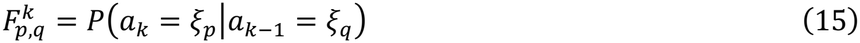

Assuming DV evolution follows a Markov process:

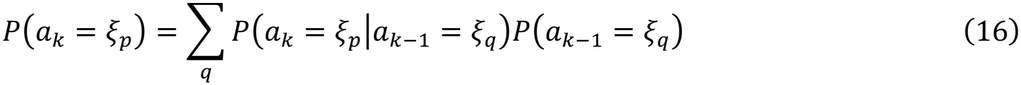

DV evolution starts at an initial value of zero, and the diffusion component follows a Wiener process. At each time step *k*, the transition probability from *a*_k−1_ to *a*_*k*_ follows a Gaussian distribution, where the mean is *µ*_*k*_ and the variance is 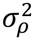 *dt*. Here, *ρ* denotes the evidence strength (*ρ ∈* {1, 0.8, 0.6}). This Gaussian distribution is given by:

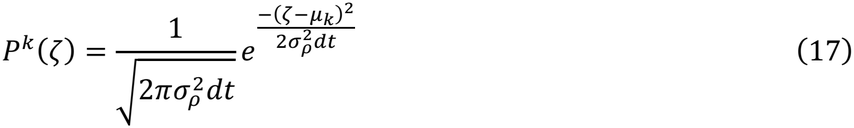

This Gaussian distribution was discretized using a finer bin size than that applied to the DV *a*, with a range extending 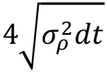 on each side of *µ*_*k*_ , covering 99.99% of the total probability mass. Under such conditions, the transition probability from *ξ*_*q*_ to *ξ*_*q*_at time step *k* is:

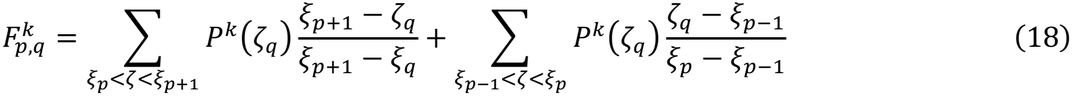

Written in matrix form, the probability distribution of DV at time step *k* follows:

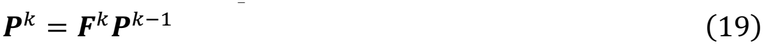

Through this iterative update, the DV distribution at stimulus offset is obtained, and the left and right choice probabilities are calculated based on equation (11). Finally, the negative log-likelihood (NLL) function is constructed for optimization.

During optimization, automatic differentiation in the JAX computing framework was used to compute NLL gradients, and the Adam optimizer was applied to minimize the NLL. The number of bins for discretized *a* was 20, and that for discretized *ζ* was 30. The time step *dt* was set to 0.125 s. Parameter search ranges were constrained as follows:

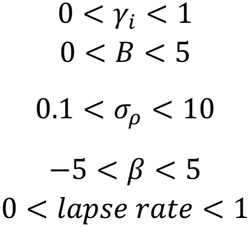

For trials with 100% evidence strength, diffusion noise was fixed at *σ*_1.0_ = 5. In total, the EDN model contained 10 free parameters.

### Language editing

The manuscript was edited for grammatical correctness and readability using DeepSeek-V4 (DeepSeek AI, Hangzhou, China). All AI-generated suggestions were manually reviewed and verified by the authors to ensure scientific accuracy and fidelity to the original meaning.

## Data availability

All source data in this paper are available from the corresponding authors.

## Code availability

All analysis codes are available from the corresponding author Xianlin Xia.

## Acknowledgments

We thank Dr. Liping Wang for help organizing marmoset behavioral training space. We thank Jinwei Pan for technical support in establishing the behavioral setup. We also thank the marmoset caring from marmoset facility of Center for Excellence in Brain Science and Intelligence Technology, Chinese Academy of Sciences, and Brain Science Infrastructure, Shenzhen. This work was funded by National Natural Science Foundation of China project (No. W2441011, B.L.), National Key R&D Program of China National Science (No. 2021YFA1101804, N.L.X.) and Technology innovation 2030 Major Program (No.2021ZD020370D/2021ZD0203704, N.L.X.).

## Author contributions

X.L.X. and N.L.X. conceived this study. X.L.X. developed the marmoset behavioral training protocol, designed the decision-making paradigms, conducted all behavioral experiments, built the drift diffusion model, analyzed all data, and interpreted all analytical results. X.L.X and N.L.X. designed the marmoset behavior system. X.L.X. wrote the manuscript, with editing by N.L.X. B.L. provided partial funding and experimental support for this work, and N.L.X. secured the majority of the funding.

## Supplementary figures

**supplemental Figure 1.**
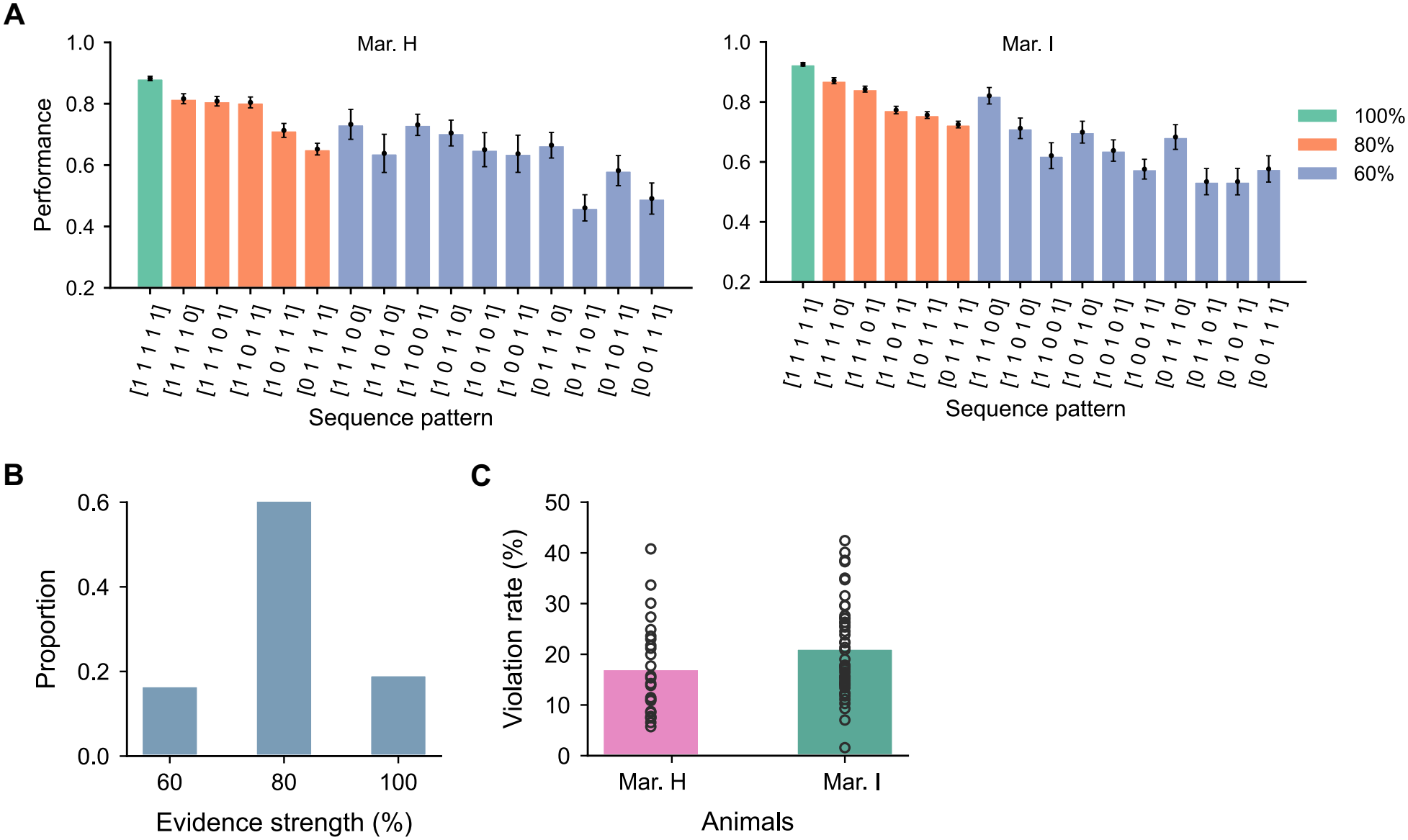
The accuracy of each sequence pattern and violation rate. (**A**) Choice accuracy of each sequence pattern (mean ± SEM) for both marmoset H (Left) and I (Right). (**B**) Distribution of trial proportions across the three evidence strength conditions (60%, 80%, 100%) in the DSTA task. (**C**) Mean violation rate of two marmosets in the DSTA task, the black circles are the violation rate of individual sessions.

**supplemental Figure 2.**
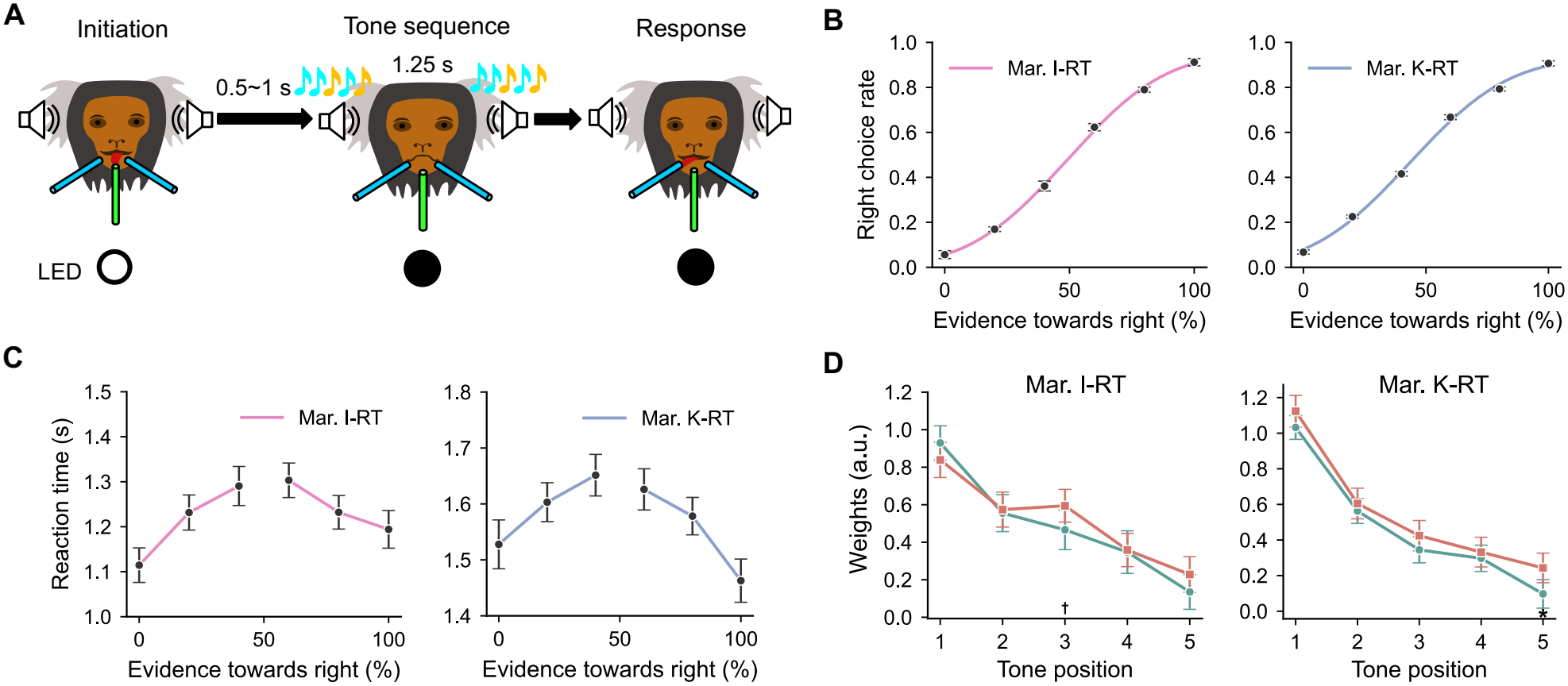
Behavioral features in reaction-time version of DSTA task. (A)Schematic of reaction-time version of the DSTA task. The marmoset can respond immediately after the stimulus onset. (**B, C**) Psychometric and chronometric curves for marmosets I and K in the reaction-time (RT) version of the DSTA task. (**D**) Comparison of temporal weighting curves for 80% (green) and 60% (orange) evidence strength (Wald test) in reaction-time version of the DSTA task. Error bars are 95% confidence interval. †: *P* < 0.1, *: *P* < 0.05, **: *P* < 0.01, ***: *P* < 0.001. For marmoset I, the *P* value for each tone position is: *P*_1_ = 0.177, *P*_2_ = 0.785, *P*_3_ = 0.0665, *P*_4_ = 0.881, *P*_5_= 0.169; For marmoset K, the *P* value for each tone position is: *P*_1_ = 0.109, *P*_2_ = 0.459, *P*_3_ = 0.163, *P*_4_ = 0.540, *P*_5_= 0.0133. The subscript numbers denote tone positions.

**supplemental Figure 3.**
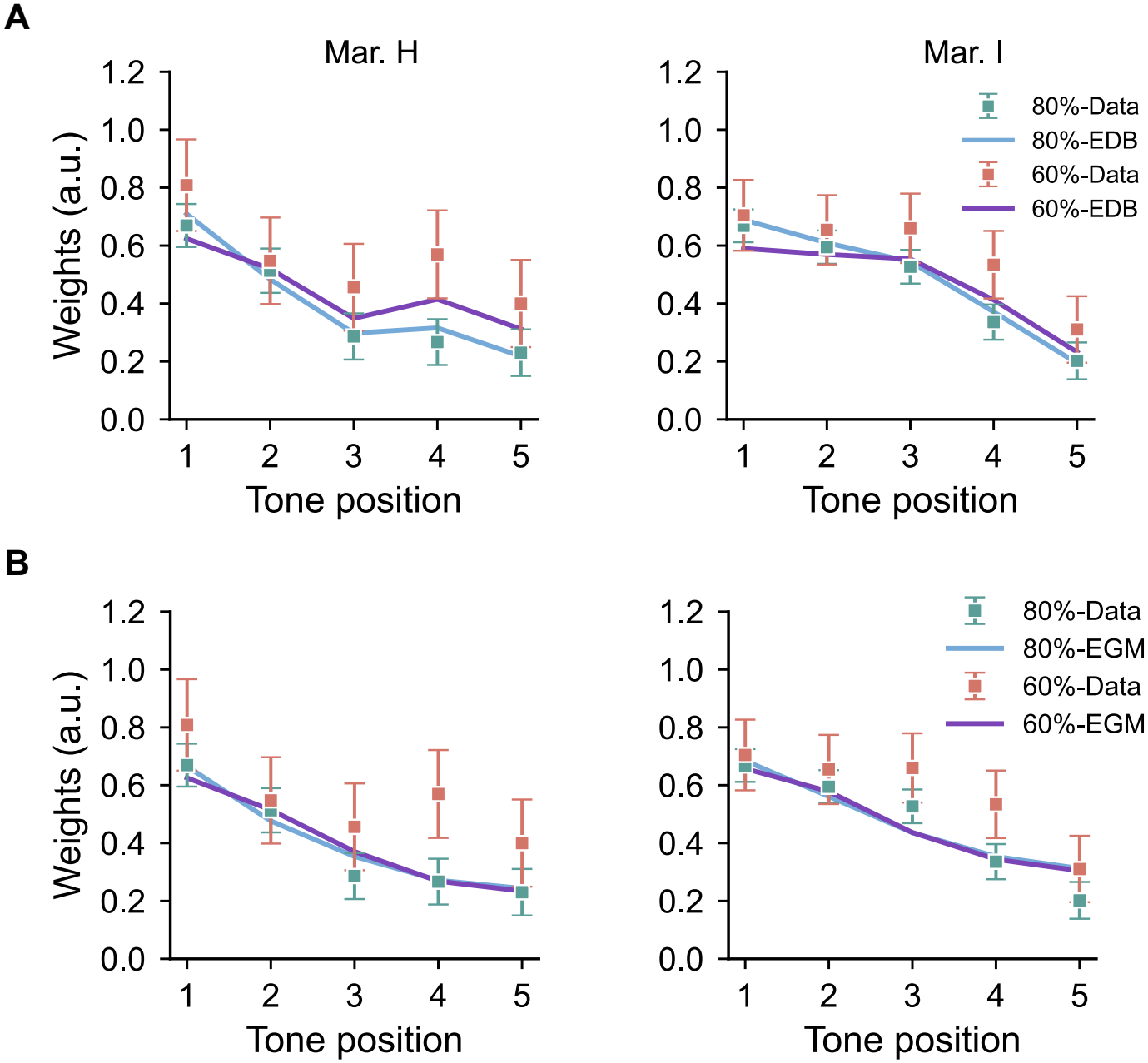
The psychophysical weights across different evidence strengths in EDB and EGM model. (**A, B**) The temporal weighting curves for DDM with evidence-strength-dependent decision bound (EDB model, panel **A**) and DDM with evidence-strength-dependent gain modulation (EGM model, panel **B**). In EDB model, the diffusion noise is set to a constant value of 5 and incorporate tone-position-dependent gain modulation into the model. For marmoset H, the *P* value for each tone position is: *P*_1_ = 0.328, *P*_2_ = 0.672, *P*_3_ = 0.555, *P*_4_ = 0.250, *P*_5_= 0.280; For marmoset I, the *P* value for each tone position is: *P*_1_ = 0.145, *P*_2_ = 0.554, *P*_3_ = 0.888, *P*_4_ = 0.531, *P*_5_= 0.570 (Wald test). In EGM model, the diffusion noise as free parameter is constant across evidence strengths and the gain modulation maintains constant across tone position. For marmoset H, the *P* value for each tone position is: *P*_1_ = 0.649, *P*_2_ = 0.655, *P*_3_ = 0.846, *P*_4_ = 0.970, *P*_5_= 0.916; For marmoset I, the *P* value for each tone position is: *P*_1_ = 0.693, *P*_2_ = 0.799, *P*_3_ = 0.993, *P*_4_ = 0.882, *P*_5_= 0.908 (Wald test). All tone positions are not significant in both models.

**supplemental Figure 4.**
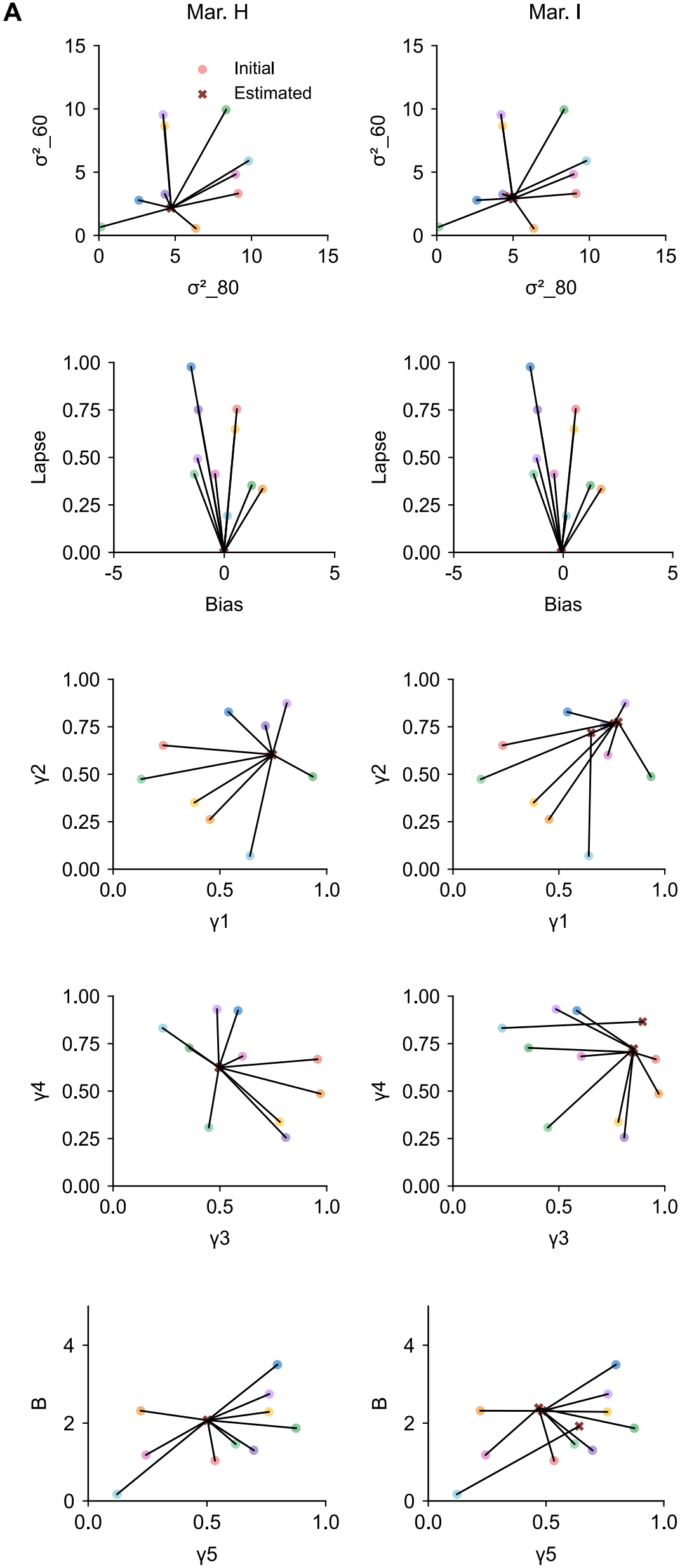
Estimated parameters with multiple random initializations. (A)Colored circles denote the values from individual random initializations. For each parameter, there are ten random initiations. Dark red crosses denote the final estimated values.

**supplemental Figure 5.**
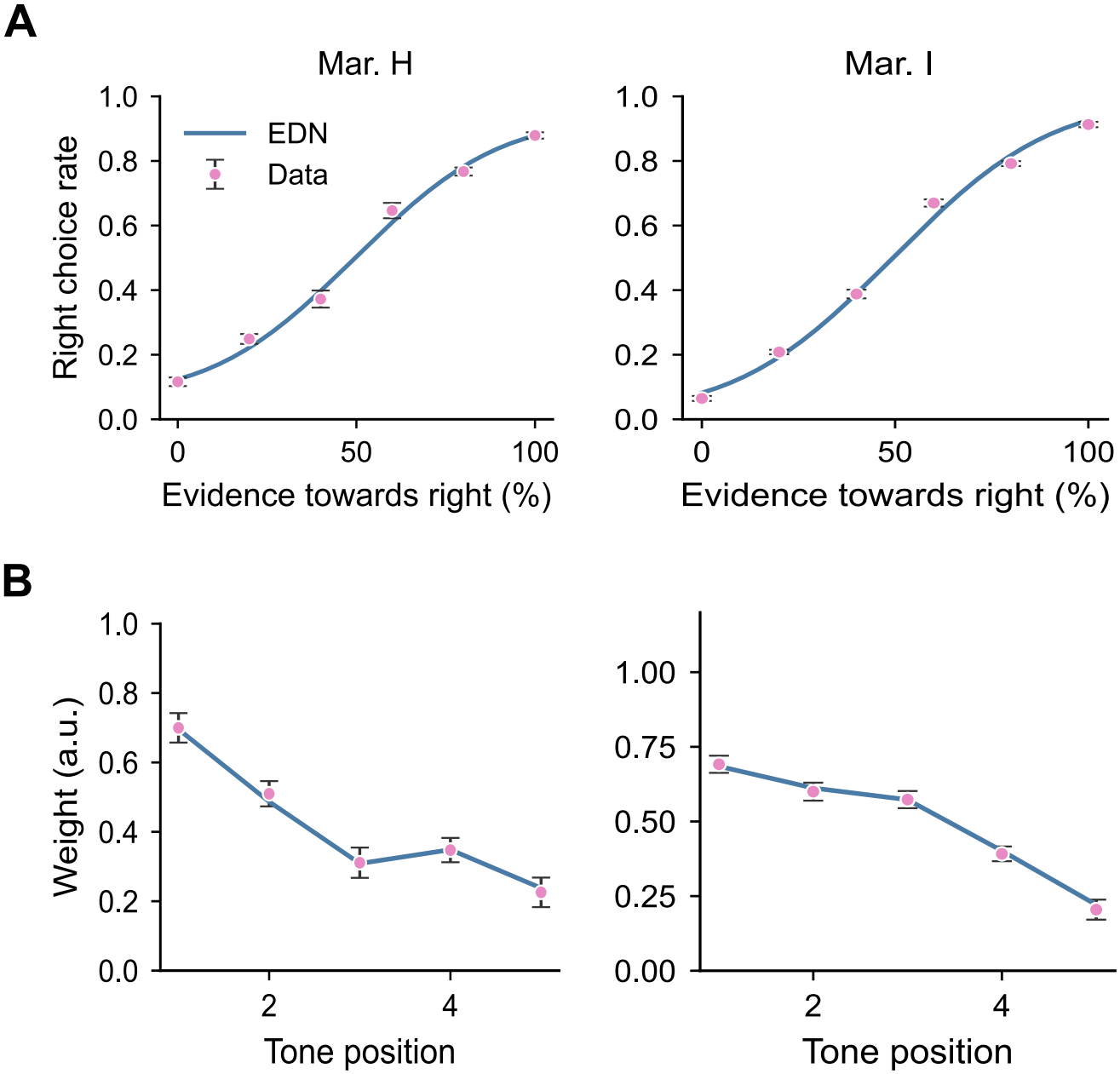
Psychometric curves and temporal weighting curves of EDN model. (**A, B**) Psychometric and temporal weighting curves for marmoset behavior (pink) and EDN model (blue).

